# Multi-scale spatial modeling of immune cell distributions enables survival prediction in primary central nervous system lymphoma

**DOI:** 10.1101/2022.09.27.509403

**Authors:** Margaretha G.M. Roemer, Tim van de Brug, Erik Bosch, Daniella Berry, Nathalie Hijmering, Phylicia Stathi, Karin Weijers, Jeannette Doorduijn, Jacoline Bromberg, Mark van de Wiel, Bauke Ylstra, Daphne de Jong, Yongsoo Kim

## Abstract

To understand the clinical significance of the tumor microenvironment (TME), it is essential to study the interactions between malignant and non-malignant cells in clinical specimens. Here, we established a computational framework for a multiplex imaging system to comprehensively characterize spatial contexts of the TME at multiple scales, including close and long-distance spatial interactions between cell type pairs. We applied this framework to a total of 1,393 multiplex imaging data newly generated from 88 primary central nervous system lymphomas with complete follow-up data and identified significant prognostic subgroups mainly shaped by the spatial context. A supervised analysis confirmed a significant contribution of spatial context in predicting patient survival. In particular, we found an opposite prognostic value of macrophage infiltration depending on its proximity to specific cell types. Altogether, we provide a comprehensive framework to analyze spatial cellular interaction that can be broadly applied to other technologies and tumor contexts.

## Introduction

The interaction between tumor cells and their surrounding non-malignant cell populations, the so-called tumor microenvironment (TME), is considered one of the major drivers in oncogenesis and tumor evolution of almost all malignancies^1^. The TME consists of a wide variety of non-malignant cells, including various classes of regulatory and effector lymphoid cells of T- and B-lineage, and various other non-haematological cell populations such as accessory cells of fibroblastic, histiocytic and dendritic nature. To adequately determine single cells in the TME within their architectural context, various cutting-edge multidimensional technologies have been developed, such as multiplex immunofluorescence (mIF) techniques (Vectra Polaris)^2,3^ and more recently higher-throughput techniques including DNA-barcoded multiplex IHC (CODEX)^4^ and imaging mass cytometry (Hyperion)^5,6^. Now that these technologies allow for accurate quantification of TME markers and record their spatial distribution, the challenge is to handle the vast amount of information that is obtained from each experiment. Therefore, advanced computational frameworks for spatial analysis are needed to translate complex cellular interplays into clinically relevant biomarkers.

Primary central nervous system lymphoma (PCNSL) is a very rare type of aggressive B-cell lymphoma that most frequently involves the brain parenchyma and more rarely the spinal cord with- or without involvement of the vitreoretinal space of the eyes^7,8^. Even though intensified treatment protocols have improved survival, 5-year survival has stagnated at only 30% which is obtained at the cost of severe medical and neuro-psychological early- and late side effects^9,10^. PCNSL has a unique underlying biology characterized by a defined genetic- and immunological context, in which immune evasion is a key mechanism^11^. On the one hand, immune evasion is mediated via genetic inactivation of antigen presentation molecules (Beta2-microglobulin, MHC class I and II) by the tumor, with subsequent loss of expression of these proteins, escaping recognition by cytotoxic CD8+ T-cells^12,13^. On the other hand, immune evasion is mediated by immune checkpoint deregulation, most frequently by translocations and chromosomal copy number alterations (CNAs) involving the *PD-L1/PD-L2* locus at 9p24.1^13^. Deregulation of the PD-1 pathway, by increased expression of PD-L1/PD-L2 leads to reversible inhibition of T-cell proliferation and activation. Studies focusing on the interactions between the malignant PCNSL cells with the intermingled immune TME currently show conflicting results^14-17^. In particular, the clinical impact of the mere quantitative presence of macrophages is inconsistent in the current literature^15,18^ and comprehensive spatial analyses are lacking. Therefore, PCNSL is a highly relevant model to explore the added value of novel computational frameworks to study the TME in detail and define its correlation with survival.

Here we propose a novel computational framework to comprehensively characterize the spatial context of the TME based on multiplex immunofluorescence (mIF). Based on marked Poisson point process theory^19^, this computational framework first captures spatial interactions between pairs of cells, such as frequent co-localization or mutual exclusiveness appears at short and/or long distance. Subsequently, downstream unsupervised- and supervised analysis were performed to identify prognostic features. We applied this framework to a set of uniformly treated PCNSL patients of the HOVON105 clinical study^20^ and identified key cellular spatial associations that distinguished PCNSL patients on outcome.

## Results

### Characteristics of the PCNSL patients from the HOVON105 study

Of 134/202 registered patients in the HOVON 105/ALLG NHL24 clinical trial^20^ biopsy material was available for pathology review and a diagnosis of PCNSL was confirmed. For the present study, for 88/134 patients biopsy material of sufficient amount and quality was selected, and processed for *PD-L1/PD-L2* fluorescence *in situ* hybridization (FISH) analysis and multiplex immunofluorescence (mIF) analysis. This patient cohort can be considered representative of the entire cohort of patients included in the clinical trial with minor, non-significant enrichment of male sex, worse performance status (WHO 3) and non-significant difference in survival (**Table 1 and Supplementary Fig. 1**).

We found frequent genetic alterations of the 9p24.1 region that contains *PD-L1/PD-L2* in tumor cells of PCNSL, determined by FISH analysis. The types of alteration include polysomy (n=3), copy number gain (n=33), amplification (n=3), and rearrangement (n=2) of the 9p24.1 region in a total of 41 out of 88 samples (47%; **Supplementary Fig. 2**). However, genetic deregulation of *PD-L1* and *PD-L2* as analyzed by FISH was not significantly associated with outcome in PCNSL patients (log-rank p-value of 0.74; **Supplementary Fig. 2E**).

### Immune-TME profiling by mIF of PCNSL cohort

Formalin-fixed, paraffin-embedded (FFPE) biopsy samples of 88 PCNSL patients were used for mIF analysis using an optimized combination of primary antibody and fluorescent dyes as indicated in Table 2. From each slide, on average 15 representative regions both within tumor areas (range 4-49, total tumor regions: 1,013) and at the interface between the infiltrating tumor border and the surrounding pre-existent cerebral tissues (range 0-17, total border regions: 380; **Fig. 1A**) were taken. For 21/88 patients, no obvious tumor border was present in the available biopsy material, and hence only tumor regions were included. Detailed analysis to determine single cells was done using InForm software (**Fig. 1**). In brief, raw mIF images (**Fig. 1B**) were segmented to identify single cells (**Fig. 1C**), and individual cells were phenotyped into the following 10 phenotypes (**Fig. 1D-E**): Tumor cells (PAX5^+^ cells) with (1) and without (2) PD-L1 expression, T-cells (CD3^+^ cells) and their subtypes classified by expression of CD8 and PD-1 (3-6), macrophages (CD163^+^ cells) with (7) and without (8) PD-L1 expression, and other cell types with (9) and without (10) PD-L1 expression (**Fig. 1B**). Cells that did not stain for any of the markers in the panel were not included in the downstream analysis.

**Fig. 1.**
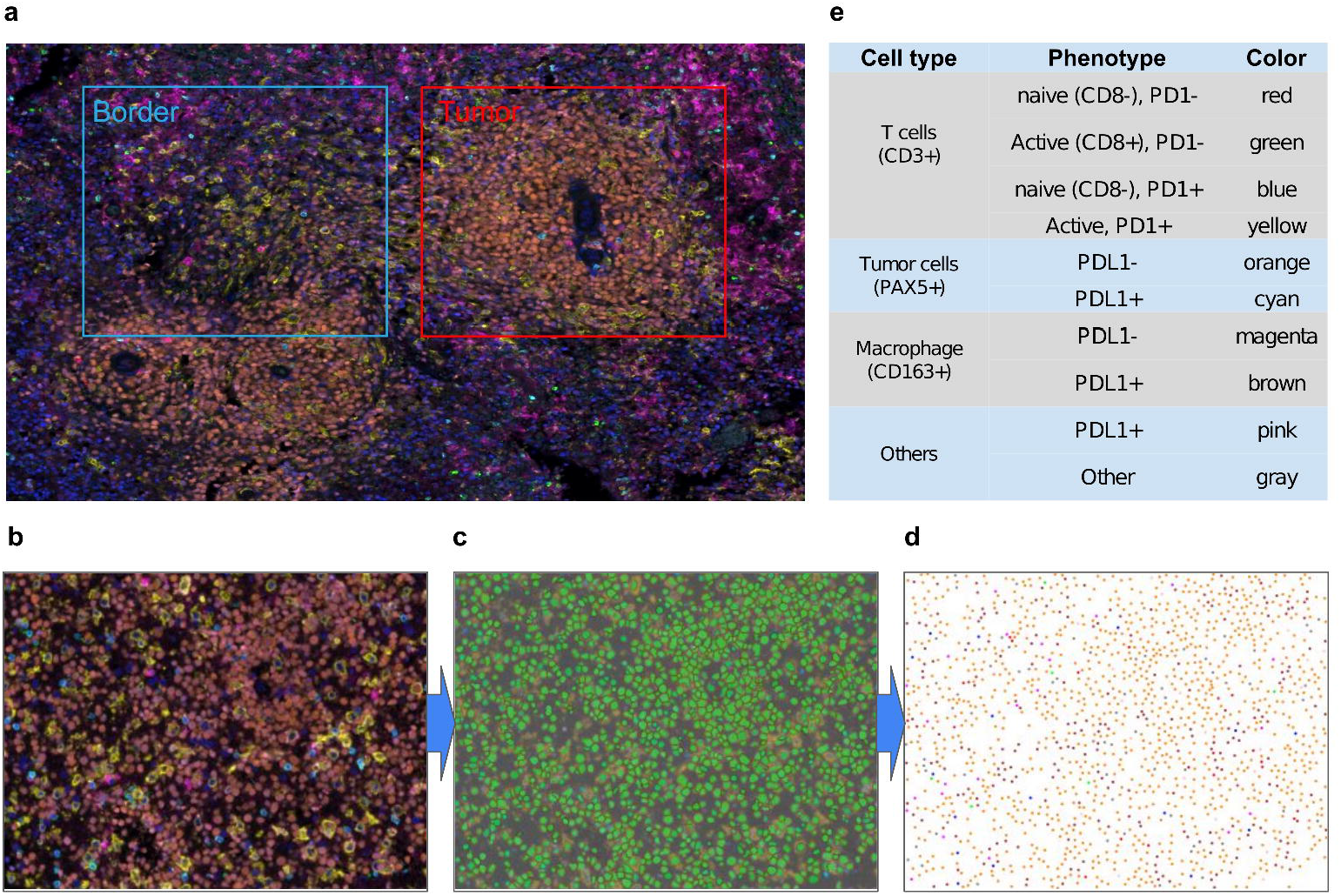
Overview of multiplex immunofluorescence (mIF) analysis of primary CNS lymphoma. **a**. For each FFPE slide, several tumor-enriched (red rectangle) and border blue rectangle) regions were selected. **b-d**. The pre-processing pipeline based on InForm takes raw mIF images (**b**), identifies cells by segmentation (**c**), and classifies cells into ten fine and coarse types (**d**), which are denoted by color defined in table (**e**; left column: cell type; middle column: phenotype based on the expression of six immune cell markers; right column: colors).

### Extraction of multi-scale TME features from mIF data

In addition to the standard non-spatial metrics such as cell counts and densities, we extracted diverse types of quantitative spatial features which collectively capture spatial characteristics of cells in PCNSL. See **Supplementary Note 1** for a detailed list of all spatial features. Among the spatial features, the statistics that capture spatial associations between two cell types are illustrated in **Fig. 2A**. We first used standard local and global spatial features that summarize spatial characteristics focusing on adjacent and all cells, including median distance between two cell types measured using only the most adjacent cells (i.e., median of minimum distances; local spatial feature) or all possible pairs of cells (global spatial feature). To further dissect spatial characteristics at multiple scales, we introduced radius-based features that describe spatial patterns in the neighborhood within a specific radius around cells. These radius-based spatial features are derived from marked Poisson point processes that enable us to calculate several statistics for 1) gaps between the cells, 2) clustering behavior of individual cell types, and 3) attraction or repulsion between two cell types. An example is Ripley’s L function which measures the average amount of one cell type within a specific radius from another cell type (**Fig. 2B**). We calculated these features based on small radii (5, 10, or 25 micrometers) and large radii (50, 75, or 100 micrometers) to cover both short- and long distance spatial interactions between cells. Representative examples of mIF images with no spatial association, attraction, and repulsion between macrophages and T-cells as captured by Ripley’s L function are presented in **Figures 2C-D**.

**Fig. 2.**
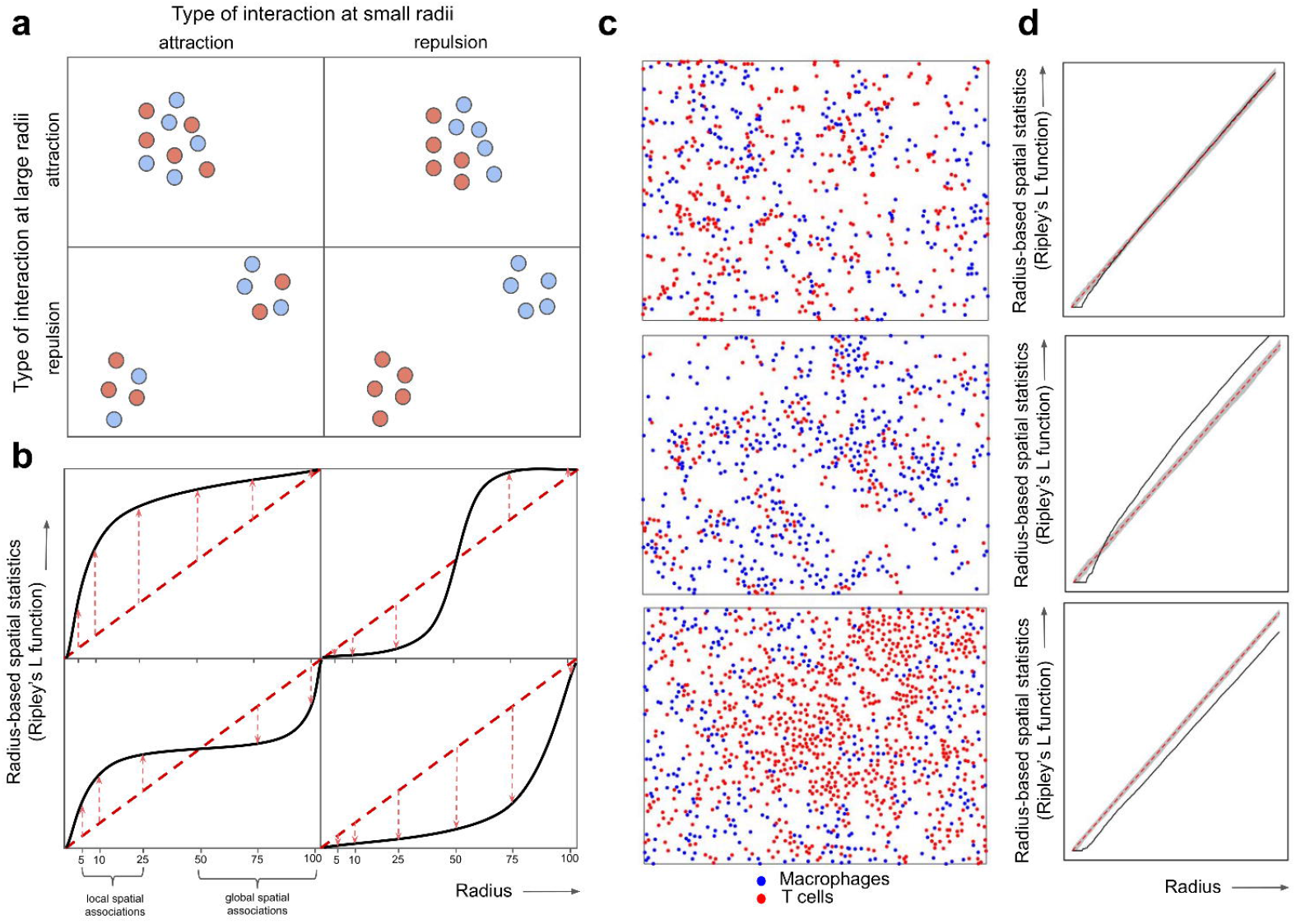
Illustration of spatial features for different types of interaction between two cell types, taking Ripley’s L function as an example. **a**. Short and long distance spatial interactions between two cell types are illustrated, categorized by attraction and repulsion at small- and large radii (denoted in columns and rows). **b**. The plots denote Ripley’s K function (y-axis) for diverse radii (x-axis) that capture short distance spatial associations (radius of 5, 10, 25 micro meters; left bottom x-axis) and long distance spatial interactions (radius of 50, 75, 100 micro meters; right bottom y-axis) for each type of spatial associations in (**a**). The dispersion of Ripley’s L function (red dotted arrows) from the theoretical values without any spatial associations (red dotted vertical line) is measured at each radius. **c-d**. Three example of processed mIF images where distinct types of spatial associations between (**c**) macrophages (blue) and T-cells (red) were captured by (**d**) Ripley’s L function.

By extracting the above-mentioned local, global, and radius-based spatial features together with the standard non-spatial features, we obtained a total of n=2,980 TME features that comprehensively characterize the TME of PCNSL patients. The high number of TME features is due to the inclusion of both coarse and fine-grind classification of total of n=13 cell types and all possible pairs among them which added up to a total n=58 cell type pairs (**Fig. 3A**). Pairing a fine-grind rare cell type (e.g., PD-1+CD8+ T-cells) with an abundant coarse cell type (e.g., macrophage) makes it less prone to missing observations than pairing it with another rare cell type (e.g., PD-L1+ macrophage). To overcome persistent missing observations, we imputed missing values by the median value per feature if the feature is observed in >50% cases (>44 samples), otherwise the observation was excluded (**Supplementary Fig. 3**). We managed to obtain more features for rare cell types, partly due to more counterpart cell types available to be paired with (**Fig. 3B**). The spatial features are more abundant than non-spatial features, particularly radius-based spatial features since they are obtained for all cell types (n=13), cell type pairs (n=58 pairs), and all six radii (**Fig. 3C**). Cumulatively, we comprehensively capture each cell type in the TME of PCNSL patients and the interplay among these cell types at multiple scales.

**Fig. 3.**
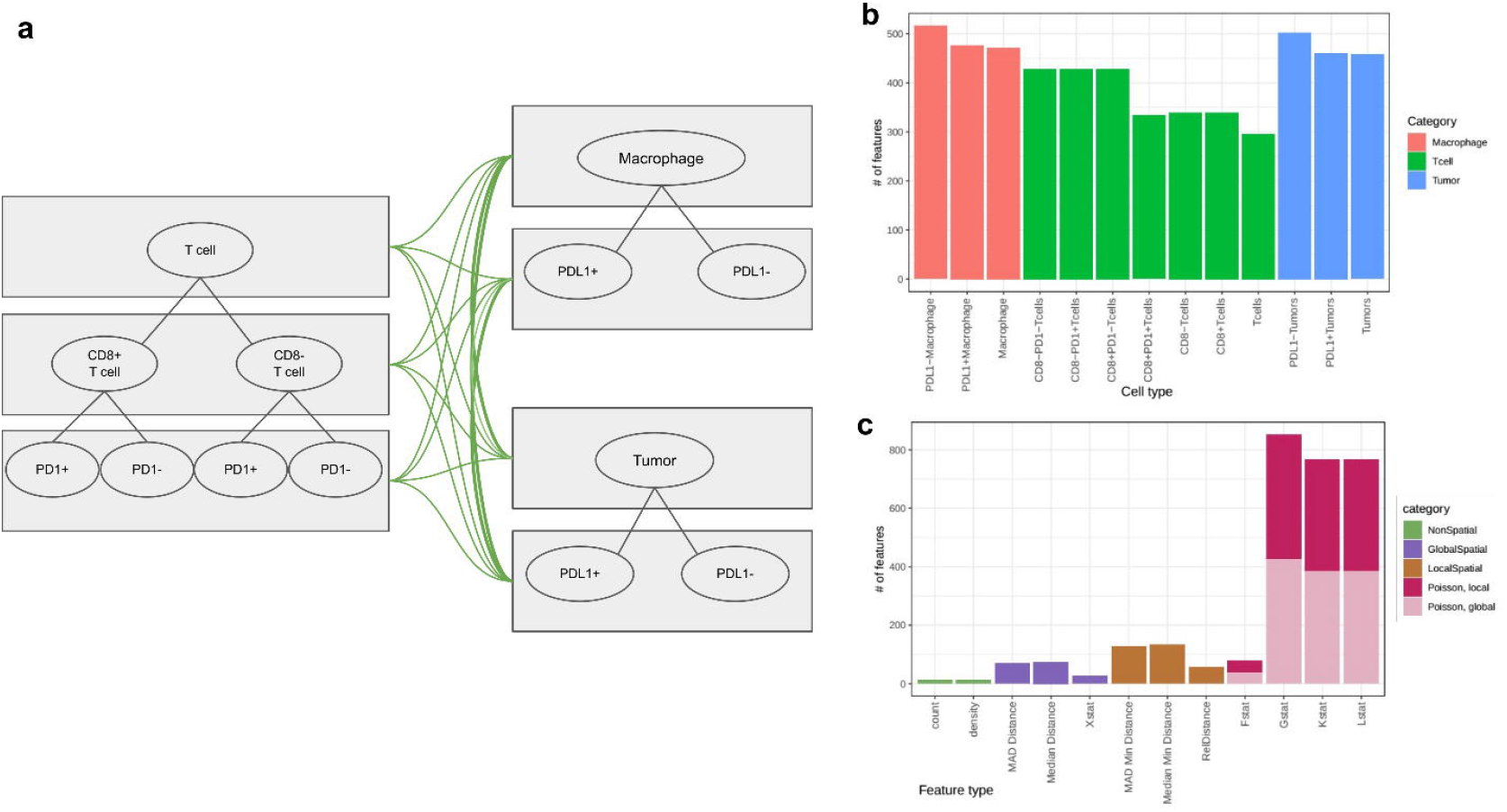
An overview of the TME features generated in this study. **a**. Pairs of cell types considered in this study. Given multiple levels of classification of T-cells (left), macrophages (top right) and tumor cells (bottom right), we considered all possible combinations of cell type pairs, denoted by green edge. **b**. The number of features per cell type in the bar plot, where colors indicate the major cell type classification. Note that the features are counted twice when two cell types are involved at the same time. **c**. Distribution of the features by their statistics in bar plots, where different types of statistics are color-coded. Radius-based spatial features are further broken down to local (r=5, 10 and 25) and global (r=50, 75, 100) features.

### Unsupervised analysis of TME features identified subgroups of patients with distinct survival outcome

To evaluate the clinical utility of the obtained TME features, we first performed clustering analysis based on cell counts only and explored the added value of spatial TME features. Unsupervised clustering analysis of immune cell counts revealed two clusters mainly separated by the presence of tumor cells, macrophages and T-cells with PD-L1 (tumor, macrophages) and PD-1 (T-cells) expression, which can further be subdivided into five clusters by the abundance of macrophages and T-cells (**Fig. 4A**). The clusters revealed a trend for survival difference (P-value of 0.064 for two clusters and 0.1 for five clusters); with a better survival in the group with higher expression of PD-1 and PD-L1 (**Fig. 4B**). However, a better prognostication could be achieved by abundance of either macrophages (p=0.0041) or T-cells (p=0.008) without accounting for the expression of PD-L1 and PD-1 (**Supplementary Fig. 4**). The clustering analysis of border regions yielded similar results with two clusters by PD-1 and PD-L1 status, but the five clusters from the border regions were not correlated with the five clusters from the tumor regions (**Supplementary Fig. 5**). There was no apparent correlation of these clusters with the outcome of the patients (**Supplementary Fig. 5**). Therefore, clustering analysis based on cell count alone, regardless of border and tumor regions, did not benefit from assessing PD-1/PD-L1 status.

**Fig. 4.**
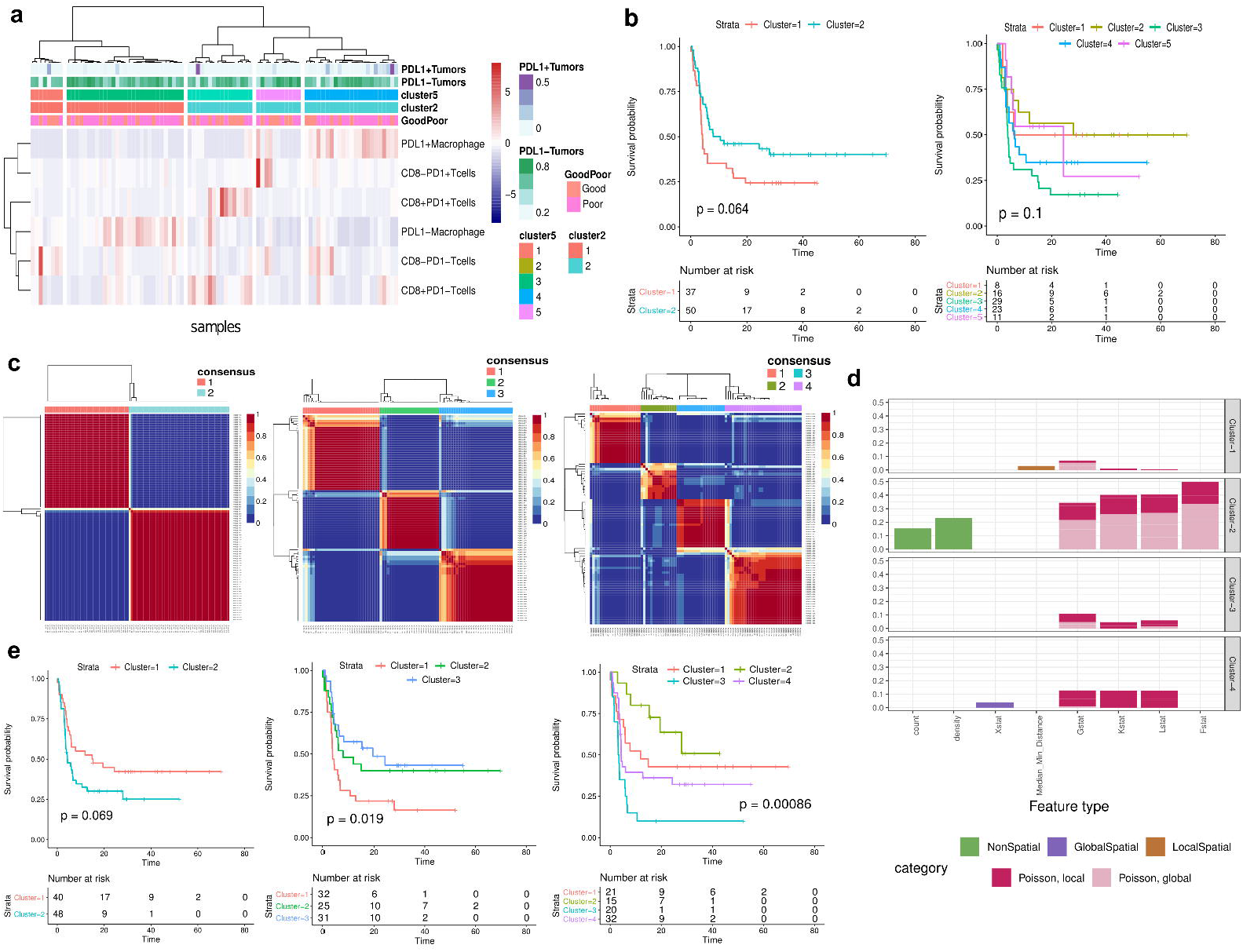
Unsupervised analysis of spatial and non-spatial TME features extracted from the PCNSL mIF data. **a**. hierarchical clustering of PCNSL samples using the counts of six non-tumor cell types. The color bars at the top provide annotations for the samples, including the normalized counts of PD-L1+ and PD-L1-tumors (first two rows), two and five clustering outcomes (third and fourth rows), and outcome classification with a 12-month threshold (bottom row). **b**. Kaplan-Meier plots comparing two (left) and five (right) clusters generated in (a). The numbers at risk are denoted below. **c**. Consensus clustering analysis of the entire TME features using NMF with a an a priori choice for different number of clusters: K=2 (left), K=3 (middle), and K=4 (right). The color denotes the frequency of NMF clustering outcome that clusters each sample pair to the same cluster. Consensus clustering outcome determined by hierarchical clustering is denoted by color bar at the top and dendrograms in rows and columns. **d**. Normalized frequency of top features (y-axis) for each of the three clusters (three panels) derived from the NMF clustering. The number of top features is normalized by the number of total features for each category and statistics. Bar colors denote the feature types. **e**. Kaplan-Meier plots comparing the survival of the subgroups derived from the NMF clustering. The number at risk and log-rank P-value are shown.

Next, we explored the impact of spatial information in addition to cell counts and detailed immunophenotypic information using non-negative matrix factorization (NMF) clustering using the entire feature set. A consensus clustering analysis with a diverse number of clusters (defined by parameter K) could identify up to four stable clusters. Less stable results were obtained as of K=5 (**Fig. 4C; Supplementary Fig. 6**). We noted that with K=2, both clusters identified non-spatial features as the top contributing feature. However, the majority of the clusters derived with a higher *K* were exclusively defined by spatial features, particularly radius-based spatial features (e.g., clusters 3-4 with K=4; **Figure 4D** and **Supplementary Fig. 7** for K=2 and K=3). Significant survival differences between the clusters were observed with clustering outcomes, particularly with higher K (p=0.069, p=0.019, p=0.00086 for K=2, K=3, and K=4, respectively; **Fig. 4E**), suggesting that combined spatial and to a lesser extent non-spatial information most adequately describes the biological impact of the TME. On the contrary, NMF clustering analysis of the border regions yielded a less stable clustering outcome with a less apparent correlation with the survival outcome (**Supplementary Fig. 8**). All in all, spatial TME features play a prominent role in identification of prognostic subgroups in PCNSL..

### Supervised analysis revealed the most predictive TME features on patient outcome

To identify which spatial TME features had the most significant impact on survival, we used supervised analysis. To this end, a Random Forest (RF) classifier was applied using the TME features from the tumor regions to predict 12-months Event Free Survival (EFS) as the clinically most meaningful dichotomizing survival cut point that is used in clinical practice. Thereby, the RF classifier achieved an out-of-bag AUC of 0.72 in models including all local, global and radius-based spatial TME features (**Fig. 5A**). The performance declined by training the same model with non-spatial features only (cell counts and densities; AUC of 0.62), further substantiating the findings on the importance of spatial features. Limiting inclusion of classes of spatial features did not improve the model (local and global spatial features only, AUC of 0.66).

**Fig. 5.**
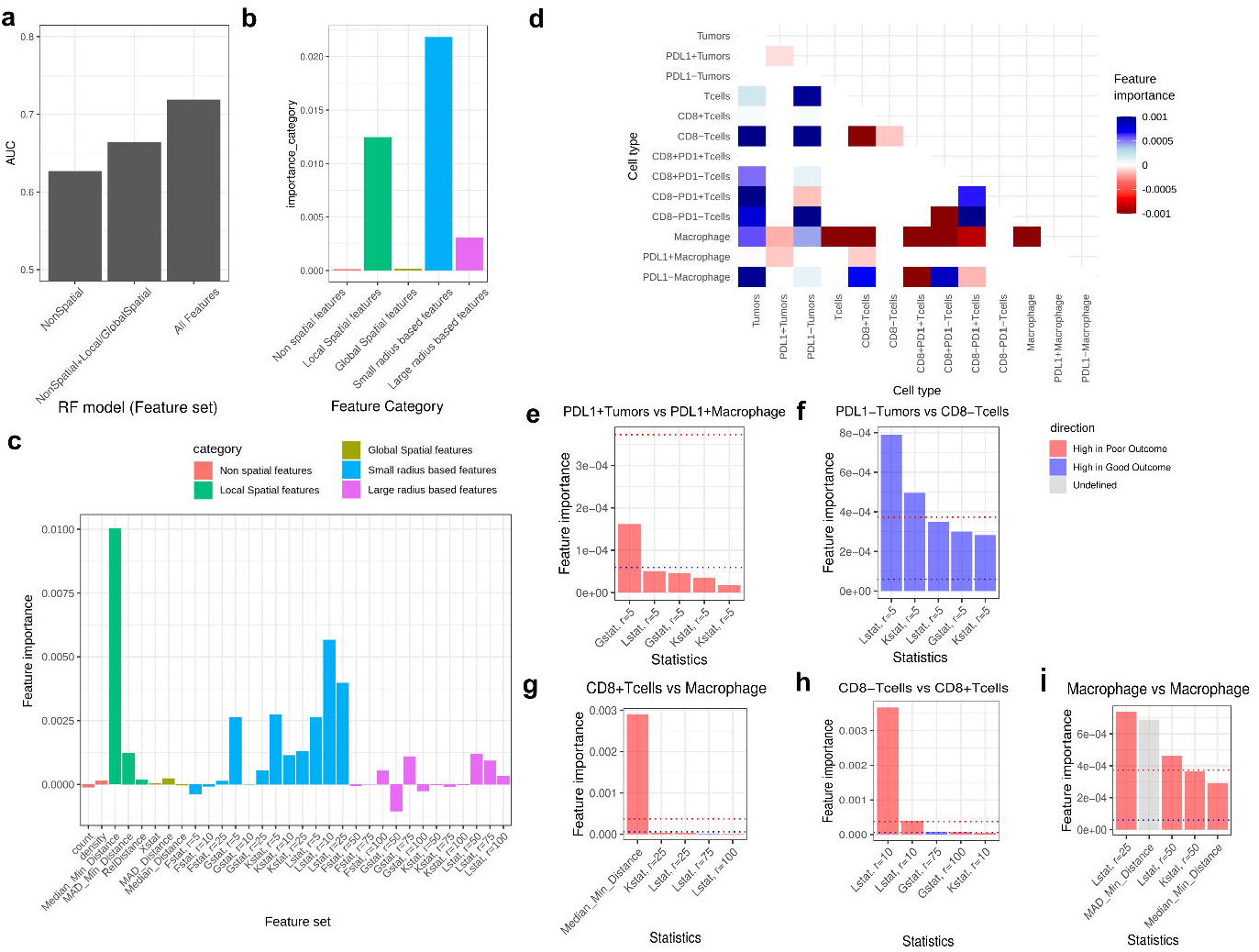
Prediction of outcome based on spatial and non-spatial TME features extracted from the PCNSL mIF data. **a**. Performance in AUC (y-axis) of RF models respectively trained by non-spatial, non-spatial combined with local/global spatial features, and entire features (x-axis). **b-c**. Importance of feature type (y-axis) in the RF model trained with the entire features classified by the type of statistics in general **(b)** and detailed **(c)** classification of feature types. The color of bars denotes the general feature type classification. **d**. Signed importance of features (color) categorized by cell types (pairs; x- and y-axis). Red and blue represent high and low in poor outcome patients, respectively. **e-i**. Importance (y-axis) of top five features in each of feature categories with the highest/lowest importance. The top feature categories by cell types (pairs) include 1) PD-L1+ tumors vs PD-L1+ macrophage (**e**); 2) PD-L1-tumors vs CD8-T-cells (**f**); 3) CD8+ T-cells vs macrophage (**g**); 4) CD8- and CD8+ T-cells (**h**); and 4) macrophages (**i**). The blue and red dotted lines in bar plots denote the 95 percentile and 99 percentile importance, and the color of bars denotes the sign of importance.

For the interpretation of the RF classifier, we next analyzed the group feature importance per feature category classified by type of statistics (**Figs. 5B-C**) and cell types (**Fig. 5D**). Among the types of statistics, radius based spatial features with a small radius up to 25 micro-meters were identified as the most important, followed by local spatial features (**Fig. 5B**). Among those features, median minimum distance was the most important, followed by L statistics with radius of 10 micro-meters (**Fig. 5C**). In general, most of the small radius-based features were important, except for the F function that measures empty space between cells. These findings indicate that close interactions between cell types are the most informative to predict patient outcome. Overall, interactions between tumor cells and non-malignant TME cell types were associated with a good outcome, except for the interaction between PD-L1+ tumors with (PD-L1 +) macrophages (**Fig. 5D**). Furthermore, it was apparent that macrophage content was associated with both good and poor outcomes, depending on its interaction partner. Patients had a poor outcome when the macrophages in the TME interact with T-cells, and a good outcome when (PD-L1-) macrophages interact with tumor cells).

From the RF classifier, we identified the top features with the highest importance for a subset of cell type pairs. First, top features for interactions between PD-L1+ tumors and PD-L1+ macrophage were mostly enriched with radius-based features with radius of 5 and are high in poor outcome patients (**Figs. 5D-E**). In the NMF clustering results with K=2 and K=4, these features were also identified as the top features in the clusters with poor survival (Cluster 2, Cluster 1, and Cluster 3 from NMF clustering with K=2, K=3, and K=4, respectively; **Supplementary Fig. 9**). The top image according to the G function with radius of 5 shows a tight interaction between PD-L1+ tumors and PD-L1+ macrophages (**Fig. 6A**). The top features for interaction between T-cells and tumors, specifically PD-L1-tumors and CD8-T-cells, reflected close interactions (radius of 5) and were high in good outcome patients (**Figs. 5D, 5F**, and **6B**). In particular, these features may explain the survival difference of Cluster 3 and Cluster 4 from NMF clustering with K=4, which are mostly derived from the poor surviving group from NMF clustering with K=2 (**Fig. 4E** and **Supplementary Fig. 9**). However, these features individually were not predictive of patient outcome (**Supplementary Fig. 10**).

**Fig. 6.**
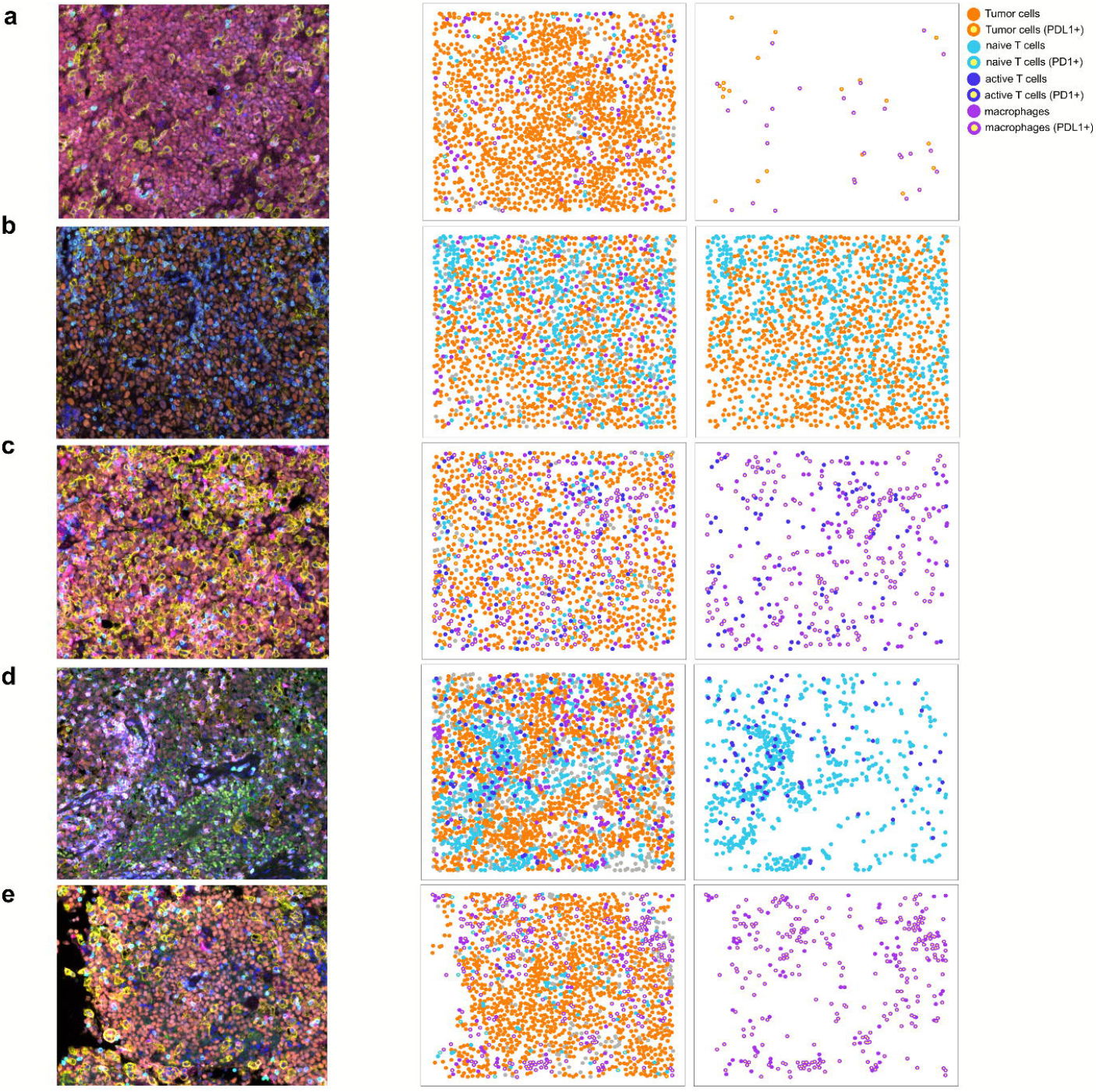
Representative images of the most important features in RF models. **a-d**. The top raw mIF images according to each of the most important features in RF (from Fig. 5e-i; left) and their processed images show all cell types (middle) and cell types (pairs) used by the corresponding feature (right). The colored dots denote individual cells and their classification, defined by each row. The cell-type classification was done differently per row according to the classification used in the corresponding feature.

Of interest, we identified important roles of the spatial context of T-cells and macrophages to predict patient outcome. First, spatial features reflecting close interactions between CD8+ T-cells and macrophages are high in poor outcome patients, among which median minimum distance being most important according to RF (**Fig. 5G** and **Fig. 6C**). Furthermore, interaction between CD8+ and CD8-T-cells as determined by the L function with a radius of 10 was associated with poor outcome (**Fig. 5H** and **Fig. 6D**). These features individually were not predictive of survival, unlike multivariate classification by RF (**Supplementary Fig. 10**). Finally, the distribution of macrophages is an important predictor for patient survival (**Figs. 5H** and **6E**). A high interaction between macrophages determined by L statistics with radius of 25, suggesting clustering of this cell type, was the most important according to RF and was significantly associated with a poor outcome (p-value of 0.00069 and FDR of 0.048; **Figs. 5H, 6E**, and **Supplementary Fig. 10**). In contrast, a similarly high abundance of macrophage is not associated with a poor survival, when their interaction is low and thus present a diffuse distribution (example in **Supplementary Fig. 11**). Taken together, a multi-scale computational framework combining frequency, architectural and spatial interaction parameters is optimal to determine the prognostic impact of TME characteristics in PCNSL.

## Discussion

We established a comprehensive computational framework based on marked Poisson point processes to characterize cellular interactions in the TME by their spatial interactions at multiple scales, encompassing general and detailed classification of cell types. Application of this framework to a well-documented cohort of PCNSL resulted in a large number of features (2,980 features extracted from 13 cell types) that subsequently required robust techniques like non-negative matrix factorization (NMF) and random forest (RF), to analyze and interpret. We identified prognostic subgroups that were specifically dictated by spatial TME characteristics rather than mere cell abundance. The high concordance between the top features from independent analysis by NMF and RF underpinned the validity of our approach. Particularly, the most important features for predicting patients’ outcomes were close spatial interactions within a radius of 25 micrometers (**Fig. 5B**), but we identified varied importance between different scales. For instance, G and K functions were the most important at a radius of 5 and 25, whereas the L function was the most important at a radius of 10 (**Fig. 5C**). This observation confirms the complementary roles of the features at diverse scales, which overall improved the performance of the RF model (**Fig. 5A**).

Previous studies on the TME of PCNSL have been mostly based on non-spatial techniques, such as transcriptome analysis^16^ or counts of single- or multiple markers by immunohistochemistry of single or few markers^17,18,21-23^. These studies suggested that a high number of PD-L1 positive macrophages may be associated with a better overall survival^16,21,22^, while the presence of tumor-associated macrophages overall rather contribute to a worse survival^15^. Furthermore, presence of perivascular and central tumor T-cell infiltrates may be predictive for a favorable outcome, although the implication of their interaction with other cell types are largely unresolved^17,23^. In the present study, we confirmed significantly worse survival in patients with a high number of macrophages and found a trend for worse prognosis in patients with a low number of T-cells. However, we showed that spatial context matters more than the abundance of these cell types. For instance, the abundance of macrophages overall was deemed a poor prognostic characteristic, but this impact did not apply in a spatial context with close spatial interaction between macrophages and T-cells. In fact, our RF model showed that the top features to predict clinical outcome were close spatial interaction of tumor cells and PD-L1+ macrophages for a good clinical outcome and close spatial interaction of macrophages and T-cells for a poor patient outcome. Importantly, we showed that close spatial interactions between different cell types were most predictive for clinical outcome in this dataset, while spatial interactions at large distances (e.g., global spatial feature and radius-based spatial features with a large radius) between cells and non-spatial features (counts and densities) were less predictive and thus not selected in the RF model. These results underscore the importance for taking spatial characteristics into account when studying the TME, not readily feasible by standard histopathological evaluation.

The sequential steps of extracting a large number of TME features at multiple scales followed by collapsing information by unsupervised/supervised analysis to reveal the key TME features with the highest biological or clinical impact may provide a basis for focused functional studies. The computational framework presented was built in a manner that is robust to handle other image-based technologies that can profile more immune cell populations, such as CODEX and mass-tag based approaches^24^. These imaging-based technologies provide an integral method to deepen our understanding on complex cellular interplays within the TME of malignancies of various nature, including malignant lymphoma.

## Methods

### Patients sample collection

Formalin-fixed paraffin-embedded (FFPE) tumor samples and clinical data were obtained from PCNSL patients enrolled in the HOVON105/ALLG NHL 24 intergroup, multicenter, open-label, randomized phase 3 study (NTR2437 and ACTRN12610000908033)^20^ through the HOVON Pathology Facility and Biobank. The pathology on all cases was reviewed according to HOVON guidelines (see **Table 1**). The HOVON 105/ALLG NHL 24 study was conducted in accordance with the Declaration of Helsinki and Good Clinical Practice, and approved by all relevant institutional review boards. Written informed consent was obtained from all patients, including use of biopsy material for research purposes.

### Fluorescence in situ hybridization for 9p24.1/ *PD-L1 /PD-L2*

Fluorescence in situ hybridization (FISH) for 9p24.1/*PD-L1/PD-L2* alterations (copy number alterations, translocations) in tumor cells was performed as previously described (Roemer et al. JCO 2016). In short, DNA of bacterial artificial chromosome (BAC) clones (Source Bioscience, Nottingham, UK) was extracted from Luria broth cultures with the Qiagen Maxi-prep kit (Hilden, Germany). The DNA was repeatedly depleted using Kreatech’s proprietary Repeat-Free technology and labeled using Kreatech’s (Leica Biosystems) proprietary ULS™ labeling. PlatinumBright™495 (green) probe targeting the 9p24.1 locus which encompassed *CD274/PD-L1*, PlatinumBright™550 (red) probe also targeting 9p24.1 and encompassing *PDCD1LG2/PD-L2* and a blue probe labeled with PlatinumBright™415 that targeted the SE9 (D9Z4) centromeric region. Approximately 100 tumor cells were analyzed for each patient manually on a Leica Biosystems DM5500B microscope, equipped with a DFC365FX camera.

### Multiplex immunofluorescence (mIF)

Multiplex immunofluorescence was performed on 4-μm-thick formalin-fixed, paraffin-embedded whole tissue sections using the Opal 7-color fluorescence immunohistochemistry (IHC) kit (Akoya biosciences, USA), as previously described^25^. In brief, slides were deparaffinized and rehydrated, followed by a blocking step for endogenous peroxidase using 0.3% H_2_O_2_/methanol and fixation with 10% neutral buffered formalin (Leica Biosystems, Germany). Slides were washed in Milli-Q water and 0.05% Tween20 in 1x Tris-Buffered Saline (TBS-T). Antigen retrieval was done by placing the slides in 0.05% ProClin300/Tris–EDTA buffer pH 9.0 in a microwave at 100% power until boiling, followed by 15 min at 30% power. Slides were cooled in Milli-Q water, washed in 1x TBS-T and blocked with Antibody Diluent (Agilent, USA). The slides were then incubated with primary antibody diluted in Normal Antibody Diluent, followed by incubation with the broad spectrum HRP from the SuperPicture Polymer Detection Kit (Life Technologies, USA). Next, the slides were incubated with Opal TSA fluorochromes diluted in an amplification buffer (Akoya biosciences, USA). The primary and secondary antibody complex was stripped by microwave treatment with 0.05% ProClin300/Tris–EDTA buffer at pH 9.0. The combination of primary antibody and fluorescent dyes is indicated in **Table 2**, in order of staining. Finally, DAPI working solution (Akoya biosciences, USA) was applied and the slides were mounted with Prolong Diamond Anti-fade mounting medium (#P36965; Life Technologies).

### Image acquisition and quantification

Stained slides were scanned using the Vectra Polaris Automated Quantitative Pathology Imaging System (Akoya biosciences, USA). From each slide, representative tumor regions and regions at the junction of tumor and surrounding cerebral tissue were selected and acquired at 40x resolution (**Fig. 1a**). After image capture, the images were spectrally unmixed (**Fig. 1b**) and analyzed, using supervised machine learning algorithms within Inform 4.2.2. (Akoya biosciences). In brief, individual cells were located and segmented to be able to perform analysis on a per-cell basis (**Fig. 1c**). Cells were then assigned into ten different phenotype categories: “PAX5+PD-L1-”, “PAX5+PD-L1+”, “CD163+PD-L1-”, “CD163+PD-L1+”, “CD3+CD8-PD-1-”, “CD3+CD8+PD-1-”, “CD3+CD8-PD-1+”, “CD3+CD8+PD-1 +”, “other PD-L1+” or “other”, based on the size of the cells and positivity of markers in the panel (**Figs. 1d-e**). In the training phase, cells representative for each category were manually selected after which the algorithm predicts the phenotype for all remaining cells. To improve accuracy, phenotypes were manually checked and the training was adjusted when necessary. This algorithm was then applied to all images of each sample. Data tables for each patient were exported for subsequent analyses.

### Feature extraction

Non-spatial features such as cell counts and densities, as well as a large number of quantitative spatial features were analyzed for 13 different cell types (**Fig. 3a**), and all possible pairs between them (n=58). For each cell type (pair) several features were calculated which represent different aspects of the spatial distribution of the cells. Marked Poisson point processes in two-dimensional space were used for these calculations, where points represent the cells and marks the phenotypes.

Local spatial features were defined by spatial patterns in terms of distances between adjacent cells. For a pair of cell types A and B, the median distance was calculated from cells of type A to the nearest cell of type B. Also, the median absolute deviation (MAD) of these distances was computed. Previously described relative distances, called SpatialScore^4^, were also included, such as the distance between tumor cells and the nearest T-cell divided by the distance between that T-cell and the nearest macrophage.

Radius based spatial features were defined by spatial patterns at a specific spatial scale, i.e. a specific radius around cells. Four distinct functions were evaluated at both small radii (5, 10 or 25 micro-meters) and large radii (50, 75 or 100 micro-meters), yielding 24 features per cell type (pair). The first function, the empty space function (F function), calculates the average amount of disk-shaped empty space between cells of a certain type. The nearest neighbor function (G function) calculates the fraction of cells of type A that have a cell of type B in the neighborhood (i.e. at distance less than a certain radius). Ripley’s K function measures the average number of cells of type B in the neighborhood of cells of type A, followed by normalization by the density of cells of type B. Ripley’s L function is a normalization of Ripley’s K function such that its theoretical value is linear in the radius. The F, G, K and L functions were normalized by their theoretical values, calculated as if the cell types were independent, homogeneous Poisson point processes.

Global spatial features were defined by spatial patterns at a global scale, i.e. how the cells are distributed over the MSI. The Chi-squared statistic measures whether the distribution of cells of a certain type is spatially homogeneous or inhomogeneous, while the median distance or MAD of distances was calculated using the distances between all pairs of cells of two cell types.

Finally, average feature values per patient for tumor and border regions were obtained for the extracted features while ignoring missing values when taking the average. Features with more than 50% cases of persisting missing values were excluded, while the rest of the missing values were imputed by the median value per feature. The feature extraction script was written in R and is available on GitHub (https://github.com/tgac-vumc/spatstat_vectra). Part of the script uses the package spatstat^19^. Details of the calculations are in **Supplementary Note 1**.

### Unsupervised analysis

Hierarchical clustering analysis was performed using the R packages *pheatmap* (v1.0.12) with Pearson correlation distance measure and Ward linkage. Consensus NMF clustering was performed using the R package *NMF* (v0.21.0) using *Brunet* update rules,with 20 iterations to determine a consensus adherence of samples per cluster and the number of clusters between 2 and 5. Top contributing features per cluster are selected by those with relative basis contribution above 0.8 compared to the maximum contributing feature (also implemented in the *NMF* R package).

### Supervised analysis

Supervised analysis using random forest (RF) was performed in R (v3.6.1) using the package randomForestSRC (v2.9.3). The number of trees was set to 100,000. Standard settings were used for the number of variables randomly selected as candidates for splitting a node (square root of number of features), and for the average number of unique data points in a terminal node (1). Predictive performance was measured by the area under the receiver operating characteristic curve (AUC) based on out-of-bag predictions. The importance of individual features and groups of features was determined by calculating (joint) permutation variable importance^26^.

## Code availability

The R script used for extracting spatial features are available at Github (https://github.com/tgac-vumc/spatstat_vectra).

## Acknowledgement

The project was supported by Dutch cancer society (KWF) grant KWF VU 2015-7925 and Leukemia research foundation, Hollis Brownstein Research grant. We acknowledge the Microscopy and Cytometry Core Facility at the Amsterdam UMC - location VUmc for providing assistance with the multiplex IHC / Vectra Polaris experiments and analyses. We also acknowledge the clinical co-principle investigator of the HOVON105/ALLG NHL 24 study Samar Issa (Department of Hematology, Middlemore Hospital, Auckland, New Zealand). We thank the collaborators at the HOVON Data Center, specifically Katerina Bakunina (HOVON Data Center, Department of Hematology, Erasmus MC Cancer Institute, Rotterdam, Netherlands) for providing excellent clinical data and support. We also would like to acknowledge Eric J. Meershoek and Danielle Hoogmoed of Leica Biosystems, Amsterdam, the Netherlands, for their invaluable contribution to the development of the 9p24.1/PD-L1/PD-L2 FISH assay and support in obtaining FISH data for this study.

