## Supplementary Materials for "Multi-scale spatial modeling of immune cell distributions enables survival prediction in primary central nervous system lymphoma"

### Multi-scale computational modeling reveals potentially prognostic immune cell interactions in primary central nervous system lymphoma

**Authors**
Margaretha G.M. Roemer[1†], Tim van de Brug[2†], Erik Bosch[2,3], Daniella Berry[1], Nathalie Hijmering[1,4], Phylicia Stathi[1], Karin Weijers[1], Jeannette Doorduijn[5], Jacoline Bromberg[6], Mark van de Wiel[2], Bauke Ylstra[1], Daphne de Jong[1‡], Yongsoo Kim[1‡*]

**Affiliations**
1. Amsterdam UMC, Vrije Universiteit Amsterdam, Department of Pathology, Cancer Center Amsterdam, Amsterdam, The Netherlands;
2. Department of Epidemiology and Data Science, Amsterdam Public Health research institute, Amsterdam UMC, Vrije Universiteit Amsterdam, Amsterdam, the Netherlands;
3. Department of Mathematics, Vrije Universiteit Amsterdam, Amsterdam, The Netherlands;
4. HOVON Pathology Facility and Biobank (HOP), Department of Pathology, Amsterdam University Medical Centre, Amsterdam, the Netherlands;
5. Department of Hematology, Erasmus MC Cancer Institute, University Medical Center Rotterdam, Rotterdam, The Netherlands;
6. Department of Neuro-Oncology, Erasmus MC Cancer Institute, Brain Tumor Center, University Medical Center Rotterdam, Rotterdam, The Netherlands;

**This PDF file includes:**

Supplementary Text
Figs. S1 to S11

**Supplementary Text**

#### Appendix: Overview of features

##### A.1 Non-spatial features

Non-spatial features count the number of cells of a specific cell type (Macrophage, PDL1- Macrophage, PDL1+ Macrophage, Tcell, CD8- Tcell, CD8+ Tcell, CD3+CD8-PD1- Tcell, CD3+CD8-PD1+ Tcell, CD3+CD8+PD1- Tcell, CD3+CD8+PD1+ Tcell, Tumor, PDL1- Tumor, PDL1+ Tumor), normalized either by the total number of cells in the MSI or by the area of the MSI.

###### A.1.1 Normalized counts

For each cell type $i$, define the normalized count by

$$\widetilde{N}_i = N_i \text{ / total number of cells in MSI}$$

where $N_i$ is the number of cells of type $i$.

###### A.1.2 Density

For each cell type $i$, define the density by

$$\lambda_i = N_i \text{ / area of MSI}$$

##### A.2 Local spatial features

Local spatial features describe spatial patterns in the data in terms of distances between adjacent cells.

###### A.2.1 Median minimal distance

Let $i, j$ be a pair of cell types (possibly $i = j$). For each cell of type $i$, calculate the distance to the nearest cell of type $j$. If $i = j$ then calculate the distance to the nearest other cell of type $i$, so that the distance is always strictly larger than 0. Let $d_{ij1}, \ldots, d_{ijN_i}$ denote these distances, where $N_i$ is the number of cells of type $i$. For each pair of cell types $i, j$, define the median minimal distance by

$$MMD_{ij} = \text{ median } d_{ij1}, \ldots, d_{ijN_i}$$

A.2.2 Median absolute deviation of minimal distances

For each pair of cell types $i, j$ (possibly $i = j$), define the median absolute deviation of minimal distances by

$$MADMD_{ij} = \text{median absolute deviation } d_{ij1}, \ldots, d_{ijN_i}$$

A.2.3 Median spatial score

For each Tumor cell, calculate the spatial score, i.e. the distance to the nearest Tcell divided by the distance from that Tcell to the Macrophage that is nearest to that Tcell. Let $s_1, \ldots, s_N$ denote these spatial scores, where $N$ is the number of Tumor cells. Define the median spatial score by

$$MSS = \text{median } s_1, \ldots, s_N$$

A.2.4 Median absolute deviation of spatial scores

Define the median absolute deviation of spatial scores by

$$MADSS = \text{ median absolute deviation } s_1, \ldots, s_N$$

**A.3 Radius based spatial features**

Radius based spatial features describe spatial patterns in the data at a specific spatial scale, i.e. a specific radius around cells. In each of the following subsections we define a function which is evaluated at 6 different radii ($r =$ 5, 10, 25, 50, 75, 100 micro-meters) yielding 6 features per function per cell type (pair). We refer to radii $r =$ 5, 10, 25 as small radii, and to radii $r =$ 50, 75, 100 as large radii.

A.3.1 Empty space function (F function)

The empty space function quantifies the amount of disk-shaped empty space between cells of a certain type. Define a grid of ?? evenly spaced points (not necessarily cells) in the MSI. For a grid point $x$ and a cell type $i$, let $d_i(x)$ be the distance from $x$ to the nearest cell of type $i$. For each cell type $i$ and radius $r$,

$$F_i(r) = \text{ mean } 1(d_i(x) \leq r)$$

where the mean is over all grid points $x$. If the cells of type $i$ are distributed as a homogeneous Poisson point process with density $\lambda_i$ and we have a very fine grid of grid points $x$, then the theoretical expected value of $F_i(r)$ equals

$$F_i^{theo}(r) = 1 - \exp(-\lambda_i \pi r^2)$$

For each cell type $i$ and radius $r$, define the empty space function by

$$\widetilde{F}_i(r) = \frac{F_i(r) - F_i^{theo}(r)}{\sigma(F_i^{theo}(r))}$$

where $\sigma(F_i^{theo}(r))$ is the theoretical standard deviation as calculated in Baddeley et al. (2015).

A.3.2 Nearest neighbor function (G function)

The nearest neighbor function is closely related to the empty space function. The difference is that the nearest neighbor function looks for disk-shaped empty space centered at a cell of a certain type, while the empty space function looks for disk-shaped empty space in general. For each pair of cell types $i, j$ (possibly $i = j$) and a point $x$ in the MSI, let $d_j(x)$ be the distance from $x$ to the nearest cell of type $j$. For each pair of cell types $i, j$ and radius $r$,

$$G_{ij}(r) = \text{mean}_i \, 1(d_j(x) \leq r)$$

where the mean is over all cells $x$ of type $i$. If $i = j$ then $d_j(x)$ is the distance from $x$ to the nearest cell of type $i$ not equal to $x$, so that $d_j(x)$ is always strictly larger than 0. If the cells of type $i$ and the cells of type $j$ are distributed as two independent Poisson point processes with densities $\lambda_i$ and $\lambda_j$, respectively, then the theoretical expected value of $G_{ij}(r)$ equals

$$G_{ij}^{theo}(r) = 1 - \exp(-\lambda_j \pi r^2)$$

For each pair of cell types $i, j$ and radius $r$, define the nearest neighbor function by

$$\widetilde{G}_{ij}(r) = \frac{G_{ij}(r) - G_{ij}^{theo}(r)}{\sigma(G_{ij}^{theo}(r))}$$

where $\sigma(G_{ij}^{theo}(r))$ is the theoretical standard deviation as calculated in Baddeley et al. (2015). In addition, we calculate $\widetilde{G}_{i*}(r)$ for each cell type $i$, radius $r$, and cell type $j$ replaced by all cells of type unequal to $i$.

A.3.3 Ripley's K function

Ripley's K function measures the average amount of cells of type $j$ in the neighborhood of (i.e. less than a certain radius from) cells of type $i$. A normalization is applied to make the value independent of the density of the cells. High values of Ripley's K function indicate attraction between cells, while low values indicate repulsion. For a pair of cell types $i, j$ (possibly $i = j$) and radius $r$,

$$K_{ij}(r) = \lambda_j^{-1} \ \mathrm{mean}_i \ N_j(B(x,r))$$

where $\lambda_j$ is the density of cells of type $j$, the mean is over all cells $x$ of type $i$, and $N_j(B(x,r))$ is the number of cells of type $j$ that are at distance less than $r$ from the cell $x$. If $i = j$ then the cell $x$ itself is excluded from the calculation of $N_j(B(x,r))$. If the cells of type $i$ and the cells of type $j$ are distributed as two independent Poisson point processes with densities $\lambda_i$ and $\lambda_j$, respectively, then the theoretical expected value of $K_{ij}(r)$ equals

$$K_{ij}^{theo}(r) = \pi r^2$$

For each pair of cell types $i, j$ and radius $r$, define Ripley's K function by

$$\widetilde{K}_{ij}(r) = \frac{K_{ij}(r) - K_{ij}^{theo}(r)}{\sigma(K_{ij}^{theo}(r))}$$

where $\sigma(K_{ij}^{theo}(r))$ is the theoretical standard deviation. In addition, we calculate $\widetilde{K}_{i*}(r)$ for each cell type $i$, radius $r$, and cell type $j$ replaced by all cells of type unequal to $i$.

A.3.4 Ripley's L function

Ripley's L function is a normalization of Ripley's K function, in such a way that the theoretical expected value is a linear function of the radius. For each pair of cell types $i, j$ (possibly $i = j$) and radius $r$,

$$L_{ij}(r) = \sqrt{K_{ij}(r)/\pi}$$

where $K_{ij}(r)$ is the unnormalized version of Ripley's K function as defined above. The theoretical expected value is

$$L_{ij}^{theo}(r) = r$$

Define Ripley's L function by

$$\widetilde{L}_{ij}(r) = \frac{L_{ij}(r) - L_{ij}^{theo}(r)}{\sigma(L_{ij}^{theo}(r))}$$

Also, $\widetilde{L}_{i*}(r)$ is calculated.

**A.4 Global spatial features**

Global spatial features describe spatial patterns in the data at a global scale, i.e. how the cells are distributed over the MSI.

A.4.1 Chi-squared statistic

The Chi-squared statistic measures how the cells are distributed over the MSI. Low values indicate a homogeneous distribution, while high values indicate an inhomogeneous pattern. Divide the MSI in 5 x 5 rectangles of equal size. For a cell type $i$ and a rectangle $R$, let $\lambda_i(R)$ be the number of cells of type $i$ in rectangle $R$ divided by the area of $R$. Define the Chi-squared statistics by

$$X_i^2 = \sum_R \frac{(\lambda_i(R) - \lambda_i)^2}{\lambda_i}$$

where $\lambda_i$ is the density of cell type $i$, and the sum is over all rectangles $R$.

A.4.2 Median distance

Let $i, j$ be a pair of cell types (possibly $i = j$). For each cell of type $i$, calculate the distance to each cell of type $j$. Let $D_{ij}$ be the $N_i \times N_j$ matrix of these distances. Define the median distance by

$$MD_{ij} = \text{median } D_{ij}$$

where the median is calculated over all entries of $D_{ij}$. If $i = j$ then the diagonal of $D_{ij}$ (which contains zeros) is excluded from the calculation.

A.4.3 Median absolute deviation of distances

Define the median absolute deviation of distances by

$$MADD_{ij} = \text{median absolute deviation } D_{ij}$$

If $i = j$ then the diagonal of $D_{ij}$ (which contains zeros) is excluded from the calculation.

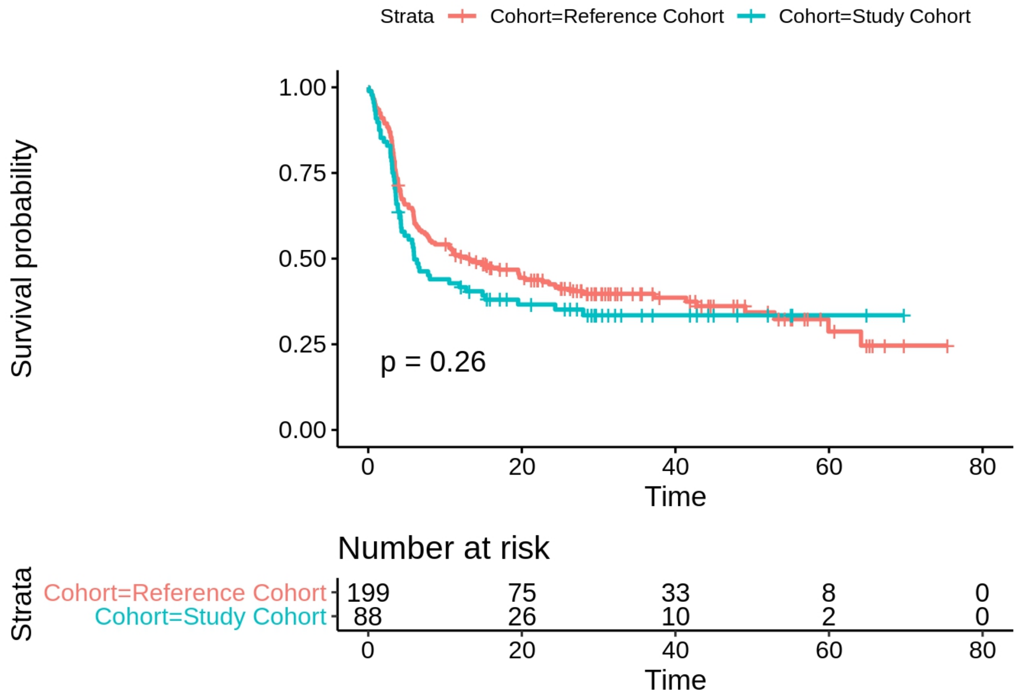


**Fig. S1. A comparison of survival outcome of reference (HOVON 105) and study cohort.** Event-free survival (y-axis) of the entire HOVON 105 cohort (red) and the subset used in this study (blue). Log-rank p-value and number at risk are indicated.

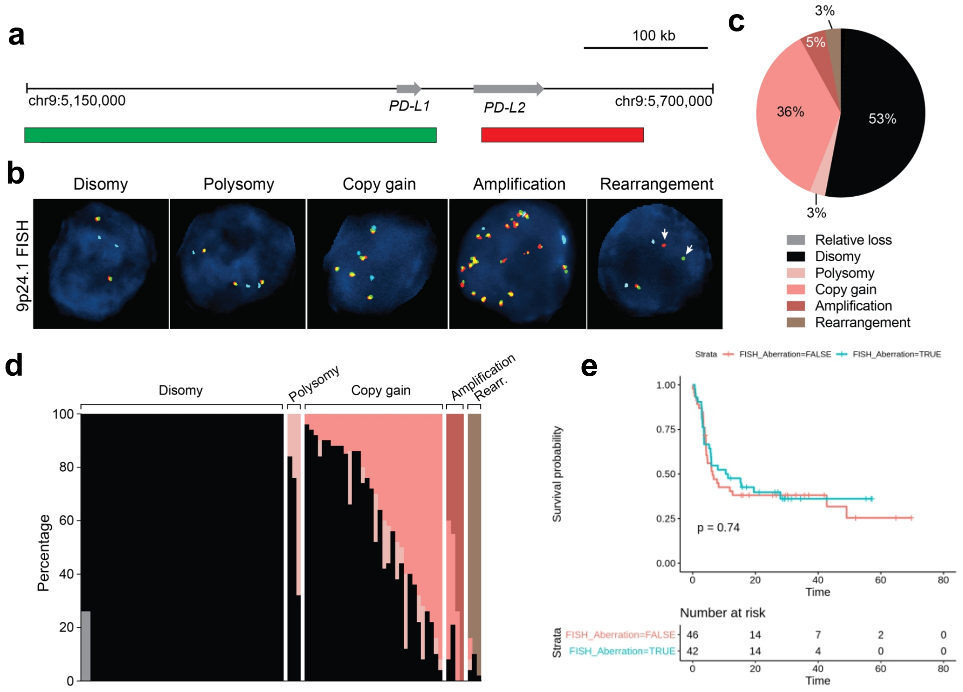


**Fig. S2. Fluorescence in situ hybridization (FISH) for the determination of CNAs and translocations of the 9p24.1/*PD-L1/PD-L2* locus. a.** A graphical representation of the target locus, indicated by green and red bars. **b.** Representative FISH images for each alteration status of 9p24.1/*PD-L1/PD-L2* locus. **c.** A pie chart represents the frequency of the 9p24.1 locus alteration status of the study cohort. **d.** A stacked bar chart represents the frequencies of cells with each 9p24.1/*PD-L1/PD-L2* alteration status in each sample, grouped by sample-wise alteration status of Disomy, Polysomy, Copy Gain, Amplification, and Rearrangement from left to right. **e.** A Kaplan-Meier plot (top) and the table for number at risk (bottom) compare event-free survival of PCNSL samples with (blue) and without (red) 9p24.1/*PD-L1/PD-L2* alteration determined by FISH. Log-rank P-value is indicated at the bottom left of the Kaplan-Meier plot.

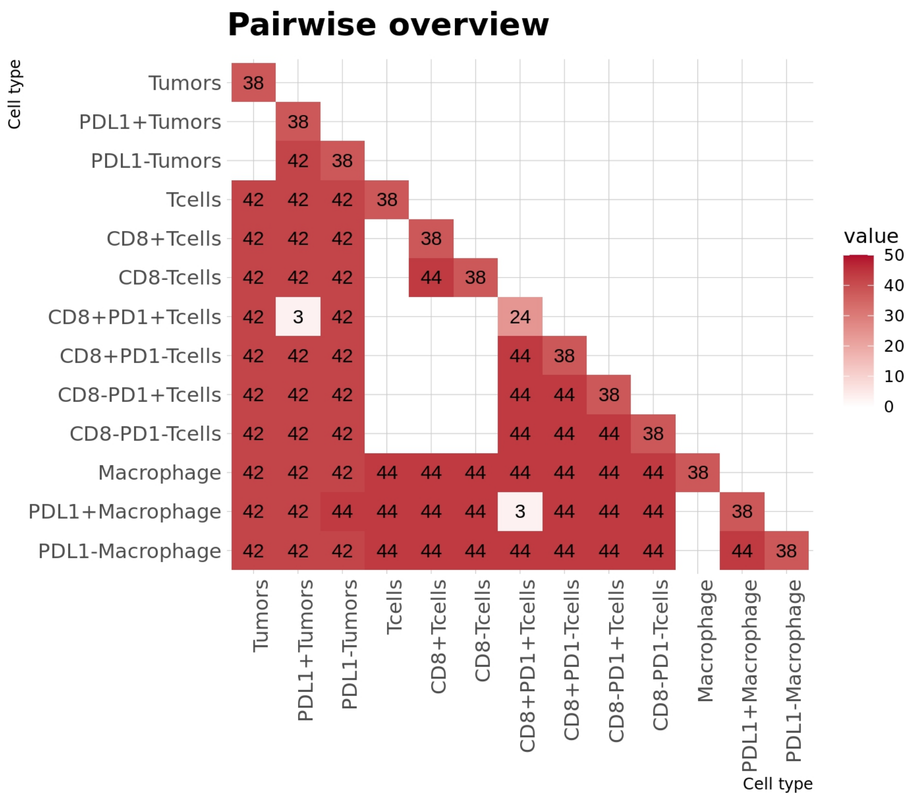


**Fig. S3. Number of features per cell type (diagonal) and per pair (lower-triangle).** The cell type pairs without the extraction of any features are left empty.

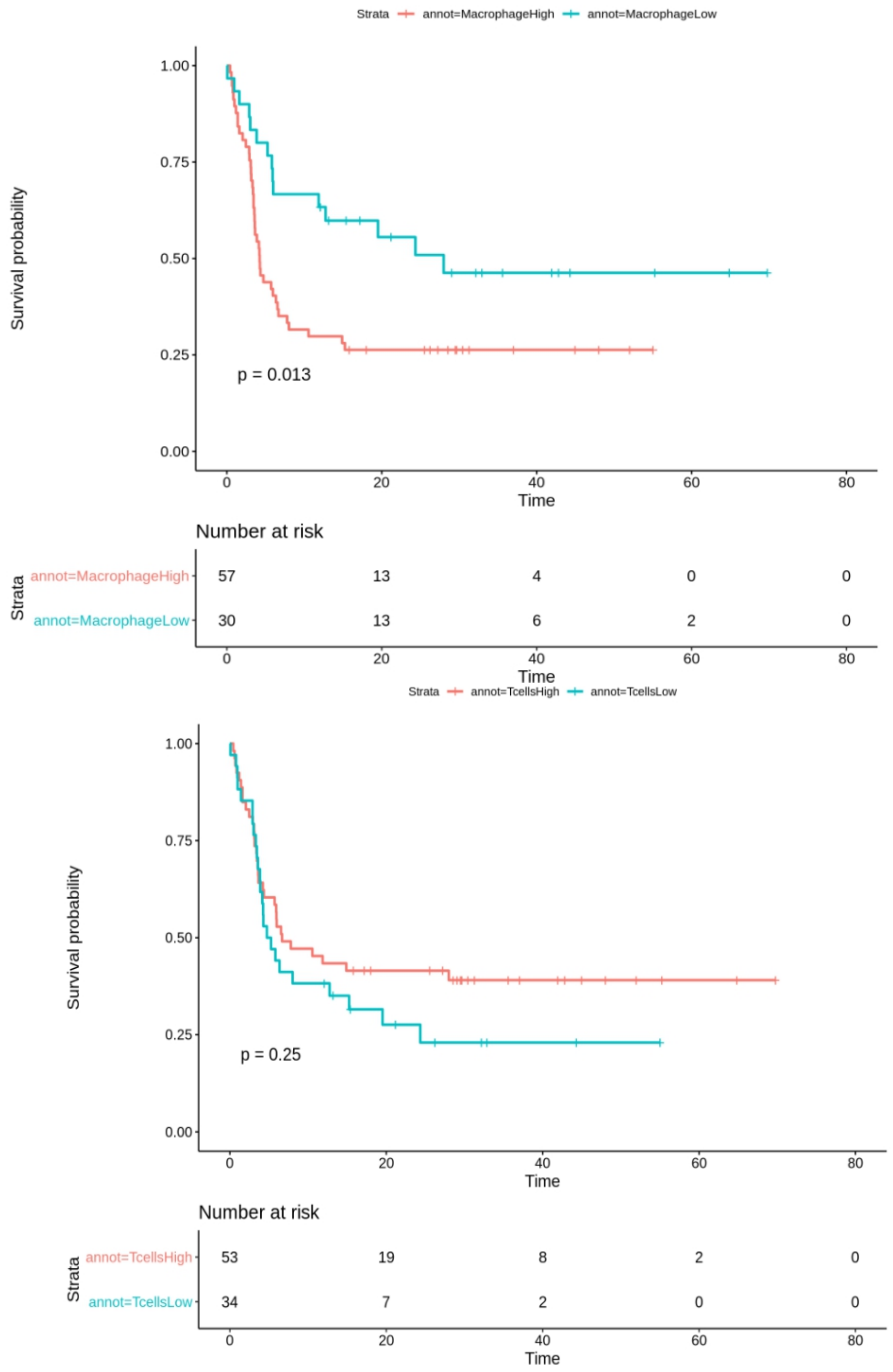


**Fig. S4. Survival difference between patients with high/low abundances of macrophages (top) and T cells (bottom).** Kaplan-Meier curves to compare the survival of patients classified into macrophage/T cell high/low based on the average cell count. Numbers at risk are shown under the Kaplan-Meier curve. Log-rank P-value is indicated at the bottom left of each Kaplan-Meier curve.

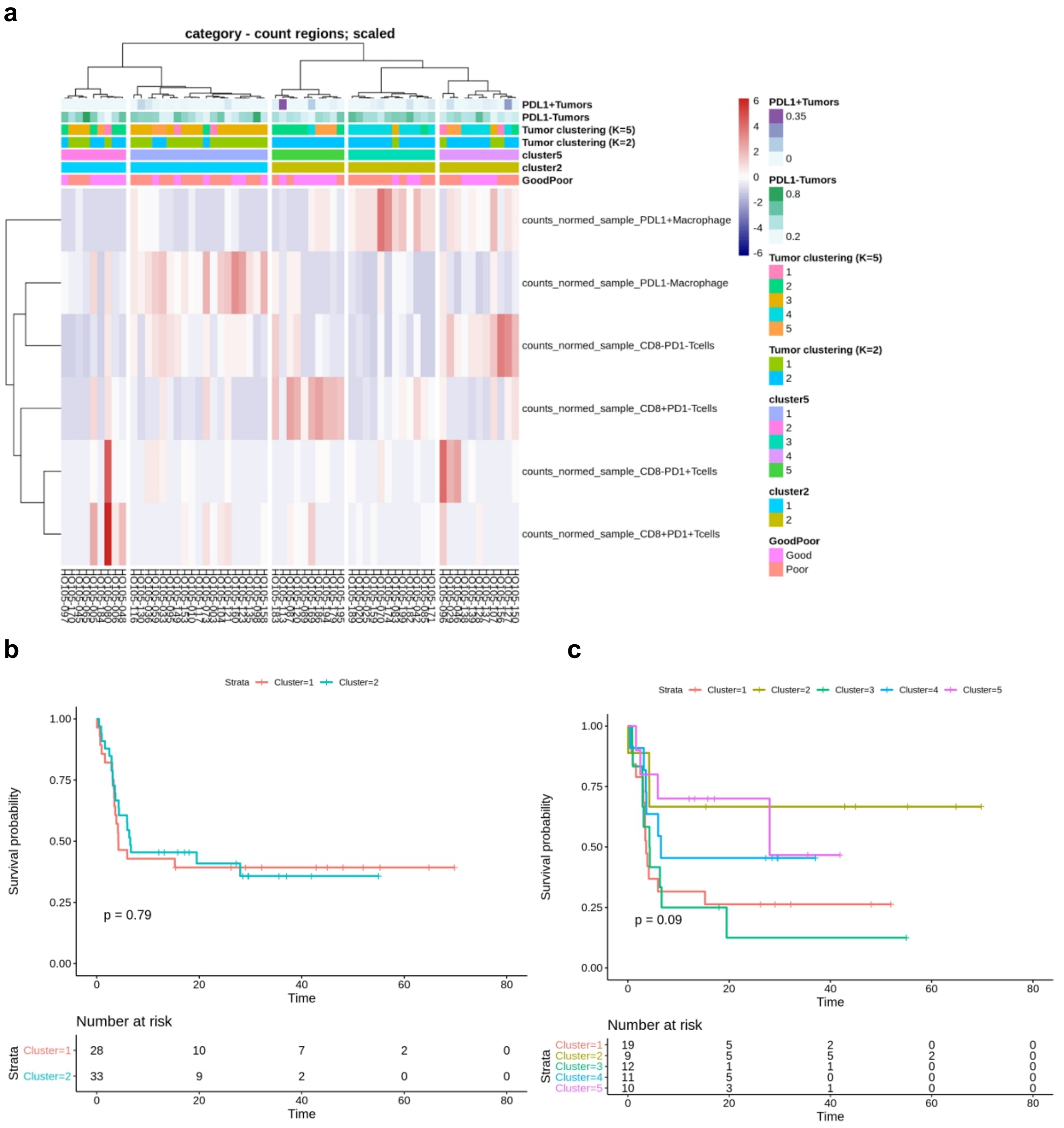


**Fig. S5. Unsupervised analysis of non-spatial TME features extracted from the border regions of PCNSL mIF data. a.** hierarchical clustering of PCNSL samples using the counts of six non-tumor cell types. The color bars at the top provides annotations for the samples, including the normalized counts of PDL1+ and PDL1- tumors (first two rows), two and five clustering outcomes from tumor images (third and fourth rows) and border images (fifth and sixth rows) and outcome classification with a 12-month threshold (bottom row). **b-c.** Kaplan-Meier plots comparing two (**b**) and five (**c**) clusters generated in (**a**). The numbers at risk are denoted below.

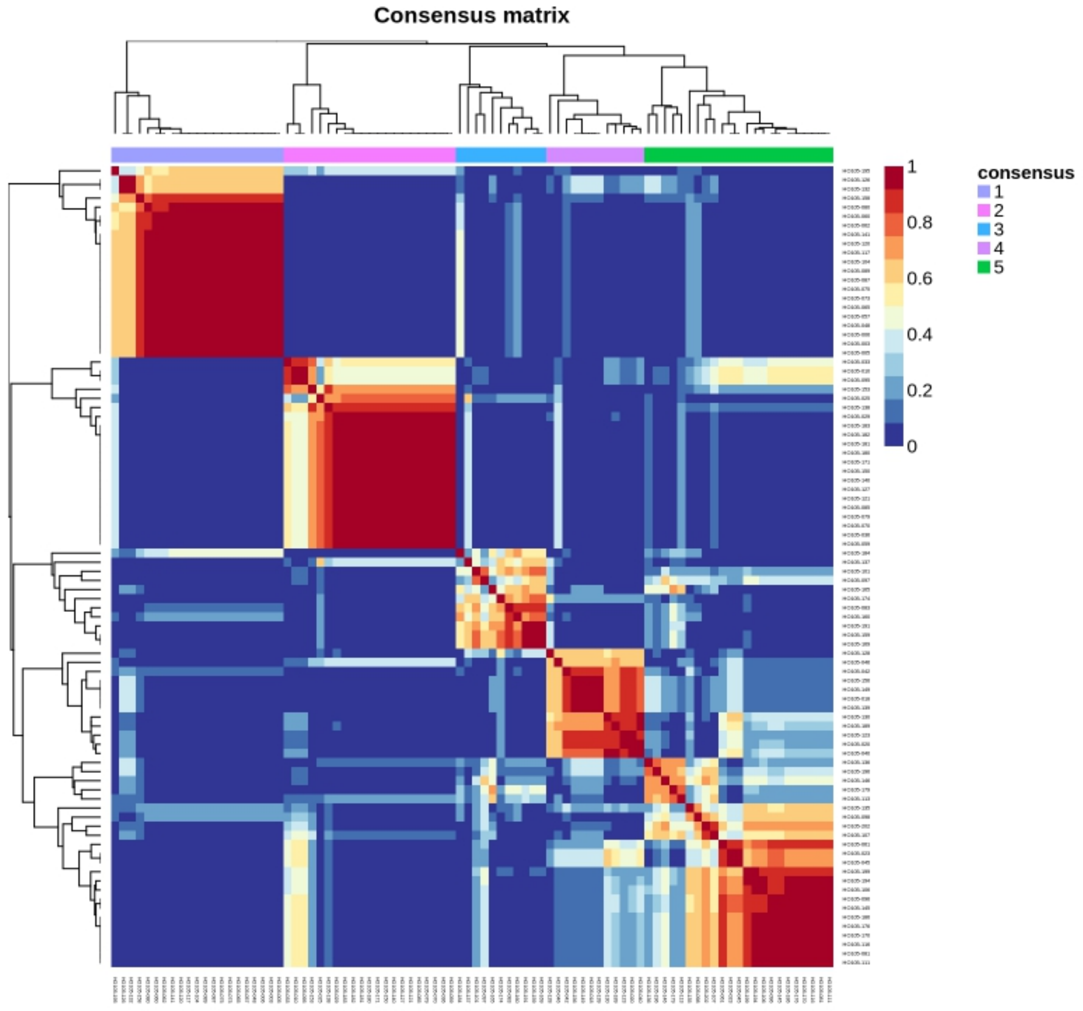


**Fig. S6. Consensus clustering analysis of the entire TME features using NMF with K=5.** The color denotes the frequency of NMF clustering outcome that clusters each sample pair to the same cluster. Consensus clustering outcome determined by hierarchical clustering is denoted by color bar at the top and dendrograms in rows and columns.

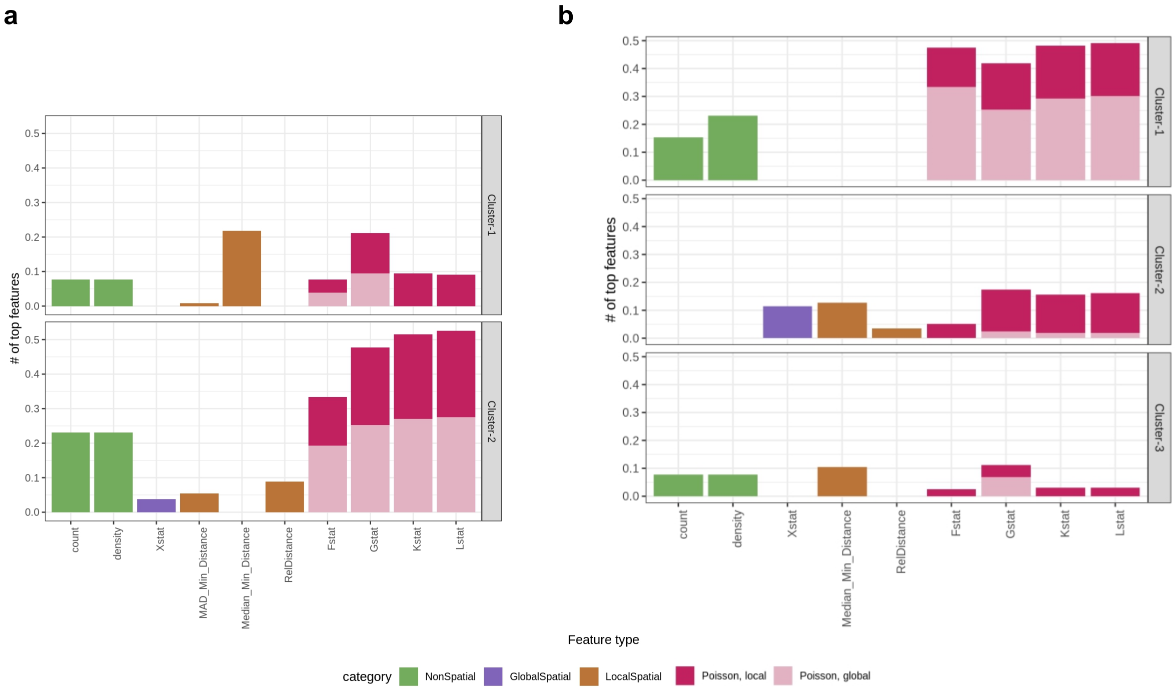


**Fig. S7. Frequency of top features of NMF clusters with K=2 and K=3.** Normalized frequency of top features (y-axis) for each of the clusters (each panels) derived from the NMF clustering with K=2 (**a**) and K=3 (**b**). The number of top features is normalized by the number of total features for each category and statistics. Bar colors denote the feature types.

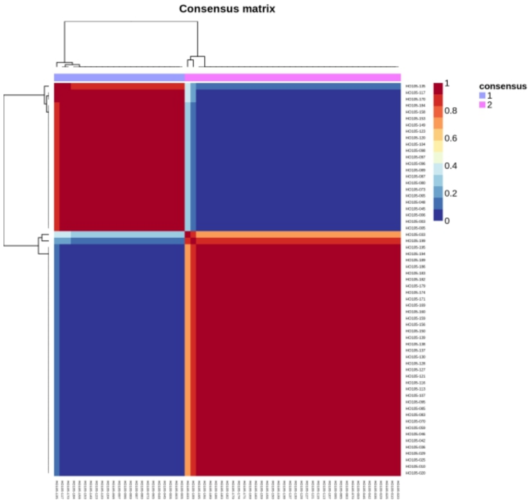
Consensus matrix
consensus
1
2
1
0.8
0.6
0.4
0.2
0

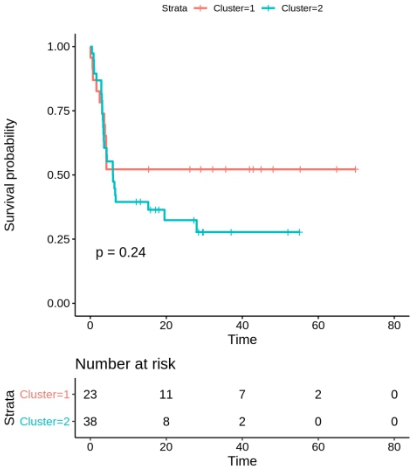
Strata
Cluster=1
Cluster=2
Survival probability
1.00
0.75
0.50
0.25
0.00
p = 0.24
Time
0
20
40
60
80
Number at risk
Strata
Cluster=1 23 11 7 2 0
Cluster=2 38 8 2 0 0

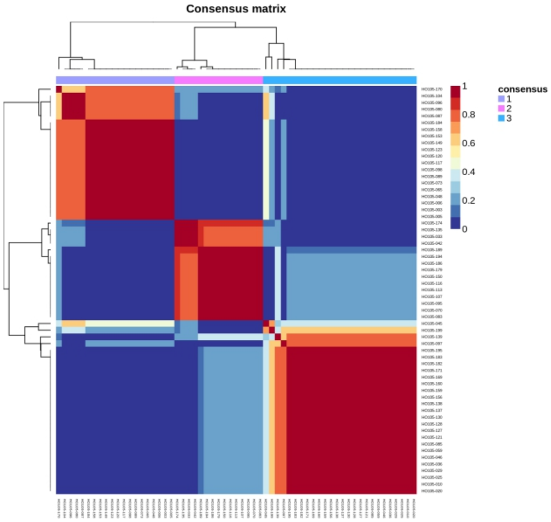
Consensus matrix
consensus
1
2
3
1
0.8
0.6
0.4
0.2
0

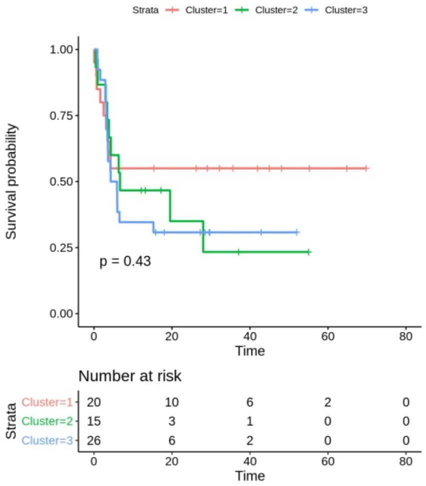
Strata
Cluster=1
Cluster=2
Cluster=3
Survival probability
1.00
0.75
0.50
0.25
0.00
p = 0.43
Time
0
20
40
60
80
Number at risk
Strata
Cluster=1 20 10 6 2 0
Cluster=2 15 3 1 0 0
Cluster=3 26 6 2 0 0

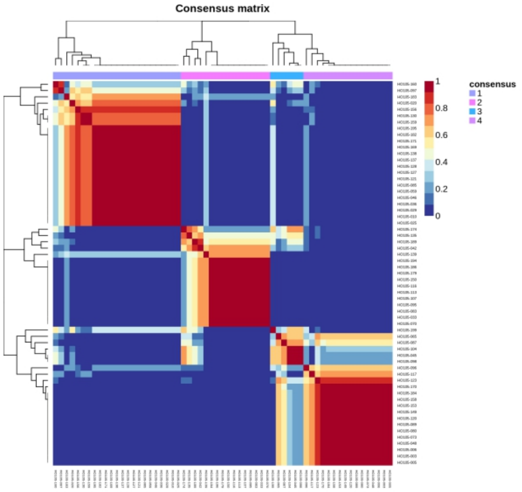
Consensus matrix
consensus
1
2
3
4
1
0.8
0.6
0.4
0.2
0

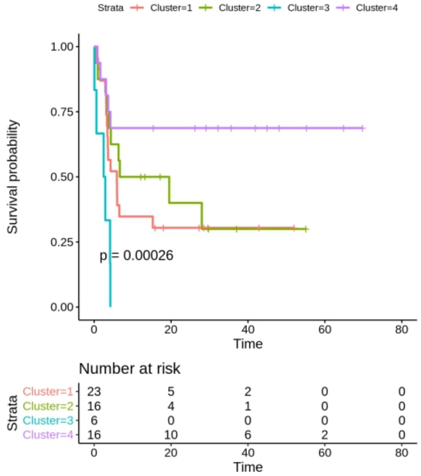
Strata
Cluster=1
Cluster=2
Cluster=3
Cluster=4
Survival probability
1.00
0.75
0.50
0.25
0.00
p = 0.00026
Time
0
20
40
60
80
Number at risk
Strata
Cluster=1 23 5 2 0 0
Cluster=2 16 4 1 0 0
Cluster=3 6 0 0 0 0
Cluster=4 16 10 6 2 0

**Fig. S8. NMF clustering analysis of spatial and non-spatial TME features extracted from the border regions of PCNSL mIF data.** Consensus clustering analysis (left) of the entire TME features using NMF with a different number of clusters: K=2 (top), K=3 (middle), and K=4 (bottom). Kaplan-Meier plots (right) compare the subgroups' survival derived from the NMF clustering. The numbers at risk are denoted under each of the Kaplan-Meier plots.

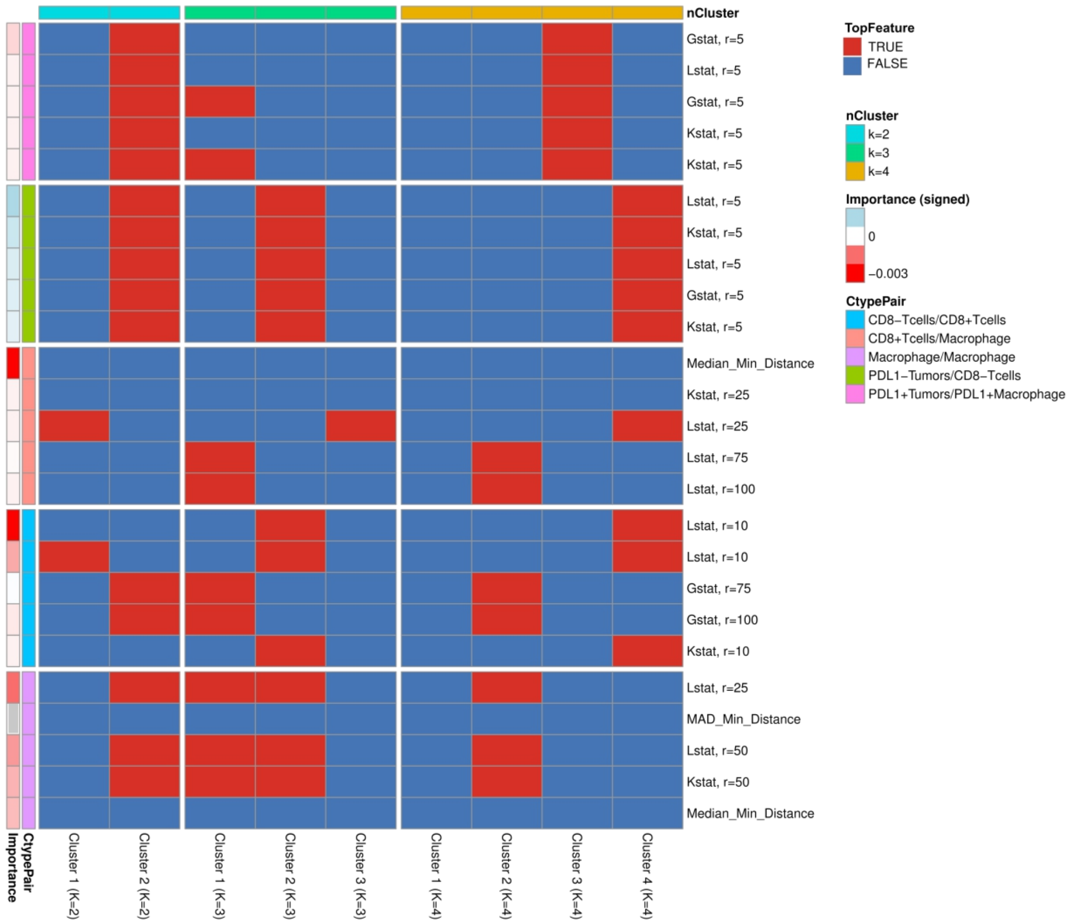


**Fig. S9. Involvement of the top spatial features from the RF in the top features for the subgroups identified by unsupervised NMF clustering.** A heatmap represents the involvement of the top features of the cell types pairs in **Fig. 5e-i** (row) among the top features that define subgroups identified by NMF clustering with K=2 (left), K=3 (middle), and K=4 (right). The signed feature importance and associated cell type pairs are indicated by color bars in the row.

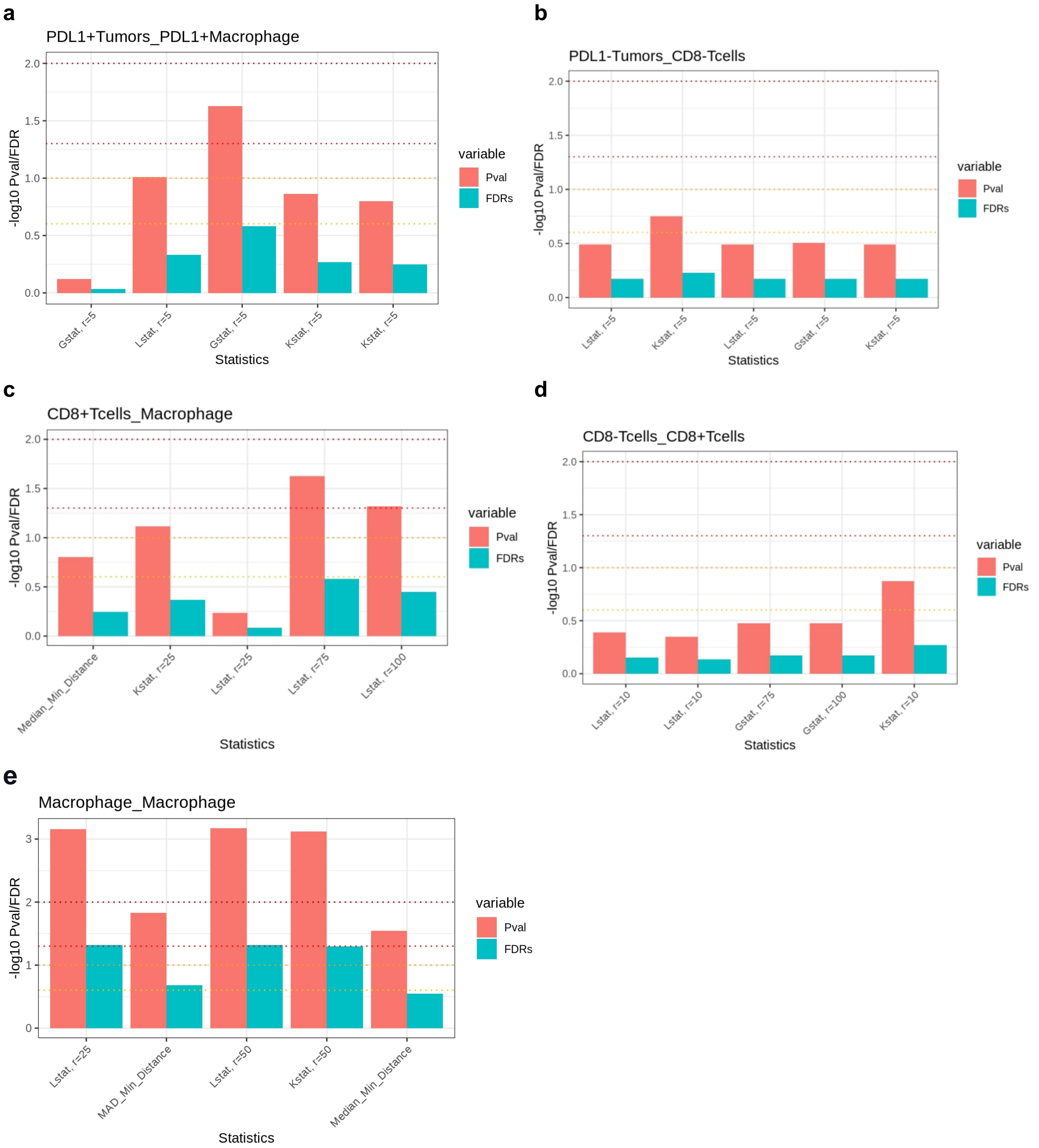


**Fig. S10. Univariate Cox regression analysis for the features with the top importance for a subset of cell type pairs.** -log10 transformed log-rank p-values (y-axis; red) and False-discovery rates (FDRs; y-axis; blue) from the univariate Cox regression analysis for the features in **Fig. 5e-i**. FDRs were obtained using the p-values from the same Cox regression for the entire 2,980 TME features. Several cutoffs (0.01, 0.05, 0.1, and 0.25) are indicated by the dotted lines (dark red, red, orange and yellow).

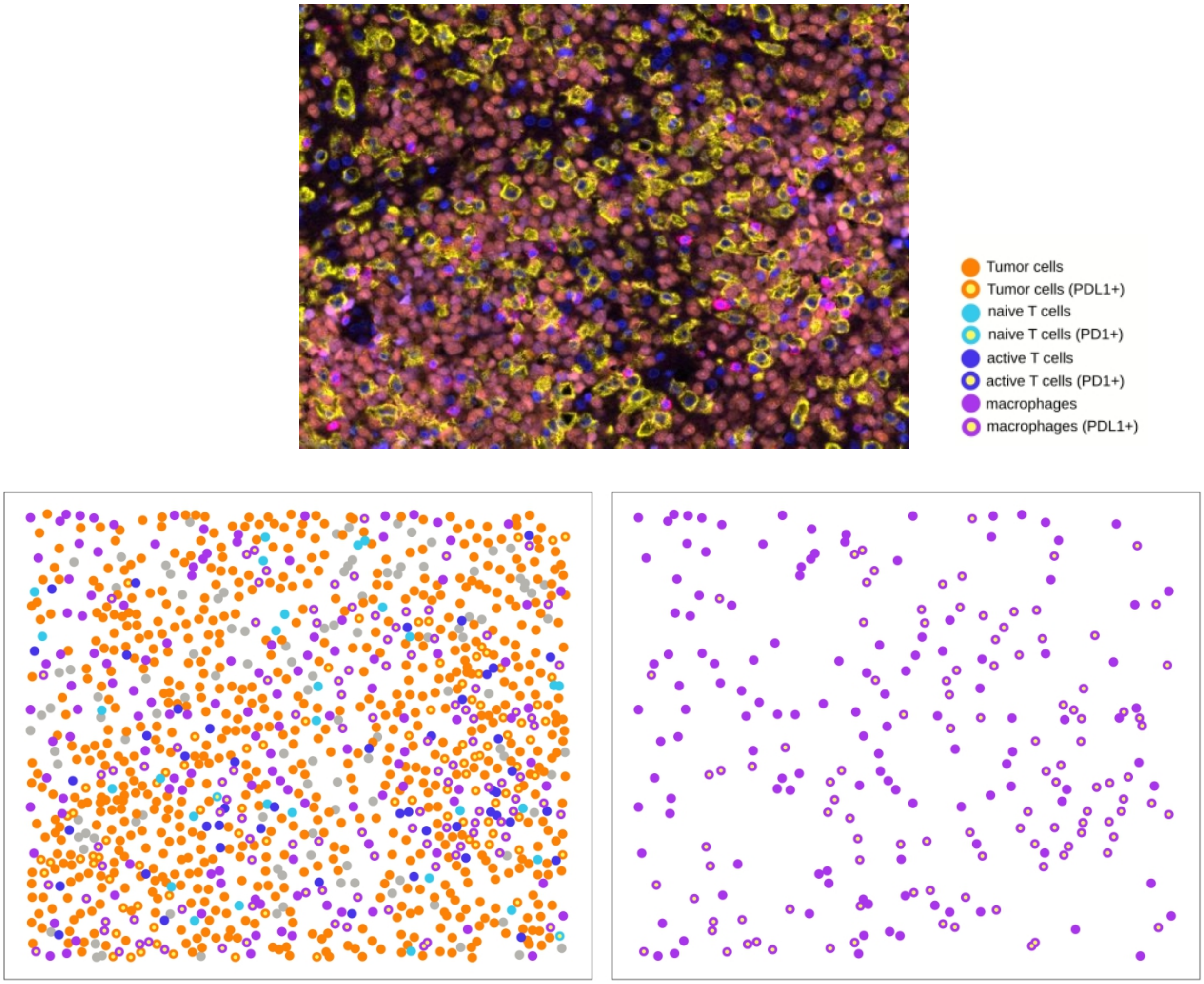


**Fig. S11. Example of mIF image with the lowest interaction between macrophage.** The raw mIF image (top) and their processed images show all cell types (bottom left) and only macrophage (bottom right). The image has the lowest macrophage interaction according to the L function with the radius of 25.
